# Adding meaning to memories: How parietal cortex combines semantic content with episodic experience

**DOI:** 10.1101/2022.10.25.513263

**Authors:** Hongmi Lee, Paul A. Keene, Sarah C. Sweigart, J. Benjamin Hutchinson, Brice A. Kuhl

**Author notes:** Correspondence to: Brice A. Kuhl. These authors contributed equally (Hongmi Lee, Paul A. Keene).

## Abstract

Neuroimaging studies of human memory have consistently found that univariate responses in parietal cortex track episodic experience with stimuli (whether stimuli are ‘old’ or ‘new’). More recently, pattern-based fMRI studies have shown that parietal cortex also carries information about the semantic content of remembered experiences. However, it is not well understood how memory-based and content-based signals are integrated within parietal cortex. Here, we used voxel-wise encoding models and a recognition memory task to predict the fMRI activity patterns evoked by complex natural scene images based on (a) the episodic history and (b) the semantic content of each image. Models were generated and compared across distinct subregions of parietal cortex and for occipitotemporal cortex. We show that parietal and occipitotemporal regions each encode memory and content information, but they differ in how they combine this information. Among parietal subregions, angular gyrus was characterized by robust and overlapping effects of memory and content. Moreover, subject-specific semantic tuning functions revealed that successful recognition shifted the amplitude of tuning functions in angular gyrus but did not change the selectivity of tuning. In other words, effects of memory and content were additive in angular gyrus. This pattern of data contrasted with occipitotemporal cortex where memory and content effects were interactive: memory effects were preferentially expressed by voxels tuned to the content of a remembered image. Collectively, these findings provide unique insight into how parietal cortex combines information about episodic memory and semantic content.

## INTRODUCTION

Neuroimaging studies have consistently implicated parietal cortex in episodic memory, leading to a number of theoretical accounts of these findings (Cabeza et al., 2008; Ritchey & Cooper, 2020; Rugg & King, 2018; Sestieri et al., 2017; Vilberg & Rugg, 2008; Wagner et al., 2005). Pattern-based fMRI studies have critically informed these accounts by showing that parietal cortex also carries information about the *content* of what is being remembered. The angular gyrus has received particular attention given that univariate activation in angular gyrus relates to the success, precision, and vividness of memory retrieval (Kuhl & Chun, 2014; Richter et al., 2016; Wagner et al., 2005) and activity patterns in angular gyrus also carry detailed information about the content of remembered events (Baldassano et al., 2017; Bonnici et al., 2016; Favila et al., 2018; Kuhl et al., 2013; Kuhl & Chun, 2014; Lee et al., 2018). However, the way in which parietal cortex *combines* memory-related and content-related information remains an important, open question (Humphreys et al., 2021; Renoult et al., 2019). In particular, at a fine-grained level, it is unclear to what degree parietal memory effects and content effects are overlapping and/or interactive. For example, are voxels that carry memory signals segregated from those that carry content information? Or, does successful remembering alter the ‘sharpness’ of content representations (Schultz et al., 2019; Woolnough et al., 2020)?

In considering the questions above, occipitotemporal cortex (OTC) serves as an important reference point. Specifically, it is well established that OTC carries robust information about stimulus content (Grill-Spector & Weiner, 2014; Haxby et al., 2001) as well as memory-related information (Martin et al., 2018; Miller et al., 1991). However, whereas memory effects in parietal cortex are typically expressed as *increases* in activation during successful remembering (repetition enhancement), memory-related effects in OTC typically manifest as *decreased* activation (repetition suppression) (Grill-Spector et al., 2006). Notably, these repetition suppression effects in OTC are thought to be stimulus-specific (Grill-Spector et al., 2006) in that they preferentially occur among voxels that are sensitive to the content of the repeated stimulus (Martin et al., 2018). This raises the question of whether a similar interaction between content and memory occurs within parietal cortex.

An important and interconnected issue is how to *measure* content representations. To date, most studies have either used (a) pattern classification algorithms that classify broad visual categories (e.g., faces vs. scenes) (Kuhl et al., 2011; Polyn et al., 2005) or (b) representational similarity analyses that test for category- or item-specific patterns of activity (Favila et al., 2018; Kuhl & Chun, 2014). While these approaches have advanced the field, they rely on a strict categorization of stimuli (whether stimuli match a category/stimulus label). This raises an important possibility that apparent content representations in parietal cortex could reflect categorization that is *induced* by task demands (Toth & Assad, 2002; Xu, 2018). This is of particular concern when experimental stimuli are deliberately selected and grouped into categories that are salient or explicitly relevant to participants (Kuhl et al., 2011, 2013; Kuhl & Chun, 2014; Polyn et al., 2005). An alternative approach is to decompose content into multiple, continuous feature dimensions and to then map these features to neural activity patterns (Huth et al., 2016; Lee & Kuhl, 2016; Pereira et al., 2018). This approach has been formalized in voxel-wise encoding models (Naselaris et al., 2011) and has been successfully applied in a handful of fMRI studies of memory to date (Bone et al., 2020; Naselaris et al., 2015).

Here, in a human fMRI study, we used voxel-wise encoding models and a novel form of content ‘tuning functions’ to test how parietal cortex and OTC combine memory signals with content representations. Specifically, using a recognition memory task with hundreds of natural scene images, we tested the degree to which memory and contents effects were spatially overlapping within parietal and OTC regions and, critically, whether recognition memory signals interacted with the expression of content information.

## METHODS

### Participants

Twelve healthy subjects were recruited from the University of Oregon community. All subjects were right-handed, native English speakers, and were in good health with no history of neurological disorder, as determined by a pre-experiment screening. All subjects reported normal or corrected to normal vision. Informed consent was obtained from all subjects according to a protocol approved by the University of Oregon Institutional Review Board, and all subjects were paid for their participation. Each subject completed two scanning sessions performed on separate days; the mean delay between sessions was five days (range: 1 – 12 days). One subject was excluded from data analysis for falling asleep in the scanner and not completing the task. Another subject was excluded for not following the instructions for the task. We therefore report results for 10 subjects (5 female) ranging in age from 18 to 29 years old (M = 23.6, SD = 3.8).

### Stimuli

The stimulus set consisted of 1284 unique color photographs of natural scenes. Images were collected from various sources on the Internet (e.g., Google Images). Content varied among the images, including people, places, animals, and objects. Included in the stimulus set were images of famous people or places; 152 (~12%) included famous people (e.g. Barak Obama), while 200 (~16%) included famous places (e.g. the Golden Gate Bridge). Each image was cropped to a 400 × 400 pixel square. 768 images were randomly selected from the image pool for each subject (384 per session). The number of ‘famous’ images was not balanced between subjects or sessions. Selected images for each session were randomly assigned to one of three experimental conditions: repeated once, repeated three times, or novel (1/3 of stimuli in each condition).

### Image annotations

Image annotations (verbal descriptions) were collected in an online experiment (**Figure 1B**) via Amazon’s Mechanical Turk using the psiTurk system (McDonnell et al., 2014). A total of 293 subjects participated for monetary compensation. Informed consent was obtained from all subjects electronically according to a protocol approved by the University of Oregon Institutional Review Board. Subjects were shown images randomly drawn from the stimulus set, one at a time. Each session had 20 unique images, except for one subject who viewed 17 images due to a technical error. For each image, subjects were instructed to type 5 to 10 words that “best represent the content or situation of the entire image.” A total of 337 sessions were completed across all subjects: 264 subjects completed 1 session, 19 completed 2, 8 completed 3, 1 completed 4, and 1 completed 7 (M = 1.15 sessions per subject, SD = 0.55). An additional 3 sessions from 2 subjects were excluded for failure to follow the instructions. While it was possible for a given subject to see the same images across different sessions, this rarely occurred: there was an average of 22.99 trials (SD = 11.07) per subject and an average of 22.94 unique images (SD = 10.74) per subject. Subjects generated on average 5.687 words per image (SD = 1.24), and each image had responses from an average of 5.23 subjects (SD = 0.81).

**Figure 1.**
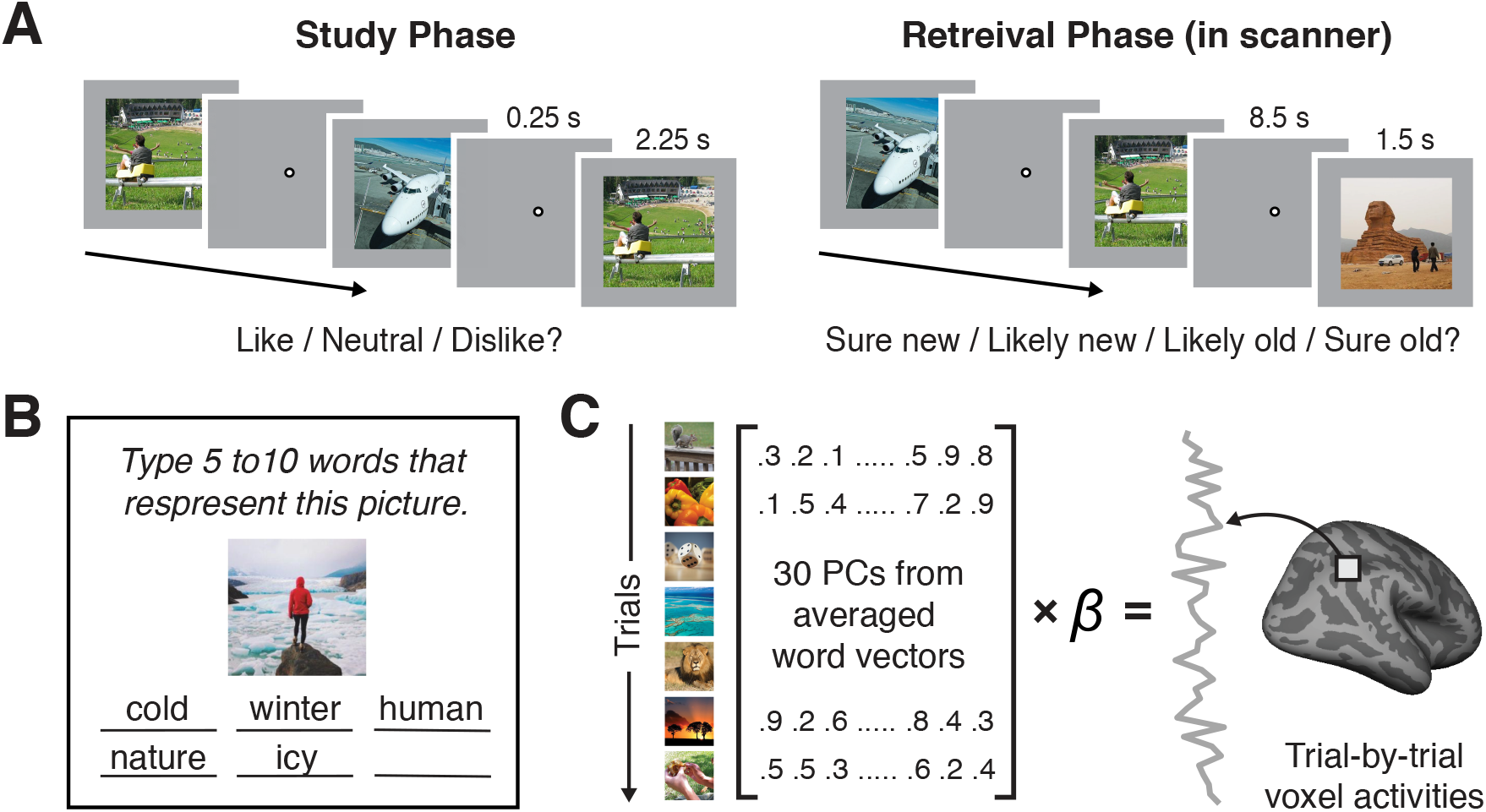
Experimental procedures and analysis methods. (**A**) Procedures for the study phase and the retrieval phase. During the study phase, subjects indicated whether they liked, disliked, or felt neutral about each presented image. During the retrieval phase, subjects performed a recognition memory test where they indicated whether they had seen each image in the study phase (‘old’) or not (‘new’), and how confident they were in their memory judgement (‘sure’ or ‘likely’). (**B**) Image annotation task. In a separate online experiment, we collected verbal descriptions of image stimuli from independent human subjects. For each image, subjects typed 5 to 10 words that best described the image. (**C**) Content-based encoding model analysis. We first performed a principal component analysis on the word embedding vectors describing the image stimuli (averaged across all words describing each image). We then used linear regression to model the relationship between each image’s first 30 principal component scores and each voxel’s activation level evoked by the image during the retrieval phase.

### Experimental design and procedures

The fMRI experiment consisted of two sessions per subject. Each session consisted of two phases: a study phase followed by a retrieval phase (**Figure 1A**). The study phase occurred outside the scanner and was intended to manipulate the episodic history of the images. During the study phase, subjects performed a pleasantness judgment task for 2/3 of the images selected for that session. Each trial consisted of an image shown at the center of the screen over a gray background for 2.25 seconds, followed by a fixation dot for 0.25 seconds. Subjects were instructed to indicate whether they liked, disliked, or felt neutral about the image by pressing a corresponding keyboard button within 2.5 seconds of the image onset. A total of 256 images were presented in the study phase for each session. Half of these images were shown only one time, and the other half were shown three times. Thus, there were 512 (128*1 + 128*3) trials in total, which were divided into 8 blocks of 64 trials each. Trial order was randomized for each session and subject. Subjects were allowed to take a short break between blocks. Subjects were told in advance that the study phase would be followed by a memory test on the items presented.

After finishing the study phase, subjects entered the fMRI scanner and completed the retrieval phase. During the retrieval phase, subjects performed a recognition memory test. Images were presented one at a time and subjects judged whether or not each image had appeared in the study phase and how confident they were in this decision. Each trial consisted of an image shown at the center of a gray screen for 1.5 seconds, followed by a fixation dot for 8.5 seconds. Subjects were asked to respond by pressing a button on the response box that corresponded to their memory judgment (sure old, likely old, likely new, sure new) within 4 seconds of the image onset. There were 8 scan runs. Each run consisted of 48 trials, divided evenly between the three experimental conditions (16 trials repeated once; 16 repeated three times; 16 novel). Stimuli in the repeated once condition had appeared once during the study phase, stimuli in the repeated three times condition had appeared three times during the study phase, and stimuli in the novel condition had not appeared in the study phase. Thus, there were 256 ‘old’ trials and 128 ‘new’ trials per retrieval session (384 trials total). Each image was shown only once during the retrieval phase, and images were not repeated across sessions. Trial order was randomized for each run, session, and subject. At the beginning and end of each retrieval session there was a 2-TR blank screen.

Sessions 1 and 2 were identical in procedure except that in Session 1 the study and retrieval phases were both preceded by practice trials. The practice trials used a separate set of images that were not included in the 1284 images used in the main experiment. There were 8 study practice trials identical to the study trials in the main experiment. There were 6 retrieval practice trials that consisted of an image presented in the center of a gray screen for 1.5 seconds, followed by a fixation dot for 3.5 seconds. Subjects could respond within 4 seconds of the image onset.

### fMRI acquisition

fMRI scanning was conducted at the Robert and Beverly Lewis Center for NeuroImaging at University of Oregon on the Siemens Skyra 3T MRI scanner. Whole-brain functional images were collected using a T2*-weighted multiband accelerated echo-planar imaging sequence (TR = 2s; TE = 25ms; flip angle = 90°; multiband acceleration factor = 2; 72 horizontal slices; grid size 104 × 104; voxel size 2 × 2 × 2mm^3^). A scanning session consisted of 8 functional runs, and a total of 244 volumes were collected for each run. Fieldmap images were also acquired to correct for B0 magnetic field inhomogeneity. A whole-brain high-resolution anatomical image was collected at the end of each scanning session using a T1-weighted MPRAGE pulse sequence (grid size 256 × 256; 176 sagittal slices; voxel size 1 × 1 × 1 mm^3^).

### fMRI data preprocessing

Preprocessing of the neuroimaging data was conducted using FSL (FMRIB Software Library, http://www.fmrib.ox.ac.uk/fsl). Functional images were first corrected for head motion within each functional run using MCFLIRT, and then across runs and sessions using linear transformation such that all functional volumes were aligned to the first volume of the first session. Motion-corrected images were then corrected for B0 magnetic field inhomogeneity using FUGUE. To more precisely coregister the unwarped images to the first volume of the first session, we performed an additional nonlinear transformation using FNIRT. Finally, functional images were spatially smoothed with a Gaussian kernel (4 mm full-width half-maximum) and high-pass filtered (cutoff = 0.01 Hz). High-resolution anatomical images were brain extracted and coregistered to the functional images using linear transformation.

### Region of interest definition

All regions of interest (ROIs; **Figure 2A**) were subject-specific and anatomically defined. For each subject, FreeSurfer’s cortical reconstruction (recon-all) was applied to the high-resolution anatomical image obtained in Session 1. Six bilateral cortical ROIs were defined based on FreeSurfer’s Destrieux atlas (Destrieux et al., 2010): In the lateral and medial parietal cortex, we examined the superior parietal cortex (SPC), intraparietal sulcus (IPS), supramarginal gyrus (SMG; a combination of the supramarginal gyrus and the Jensen sulcus), angular gyrus (ANG), and posterior medial cortex (PMC; a combination of the precuneus, subparietal sulcus, and dorsal posterior cingulate gyrus). We also examined the occipitotemporal cortex (OTC), which consisted of several brain areas spanning the occipital and ventral temporal cortex (the occipital pole, inferior occipital gyrus and sulcus, middle occipital gyrus, superior occipital gyrus, cuneus, lingual gyrus, fusiform gyrus, parahippocampal gyrus, calcarine sulcus, anterior and posterior transverse collateral sulcus, middle occipital sulcus and lunatus sulcus, superior occipital sulcus and transverse occipital sulcus, anterior occipital sulcus and preoccipital notch, lateral and medial occipitotemporal sulcus, parieto-occipital sulcus). All ROIs were masked by subject-specific whole-brain masks generated from functional images to exclude areas where signal dropout occurred. The number of voxels included in the ROIs varied across subjects (SPC range = 1707–2798, M = 2116.7, SD = 315; IPS range = 2100–3114, M = 2570.2, SD = 345.8; SMG range = 2250–3171, M = 2643.3, SD = 298.4; ANG range = 1965–2720, M = 2226.5, SD = 255.2; PMC range = 2617–4387, M = 3540.1, SD = 540.6; OTC range = 18609–24022, M = 21222, SD = 1733.9).

**Figure 2.**
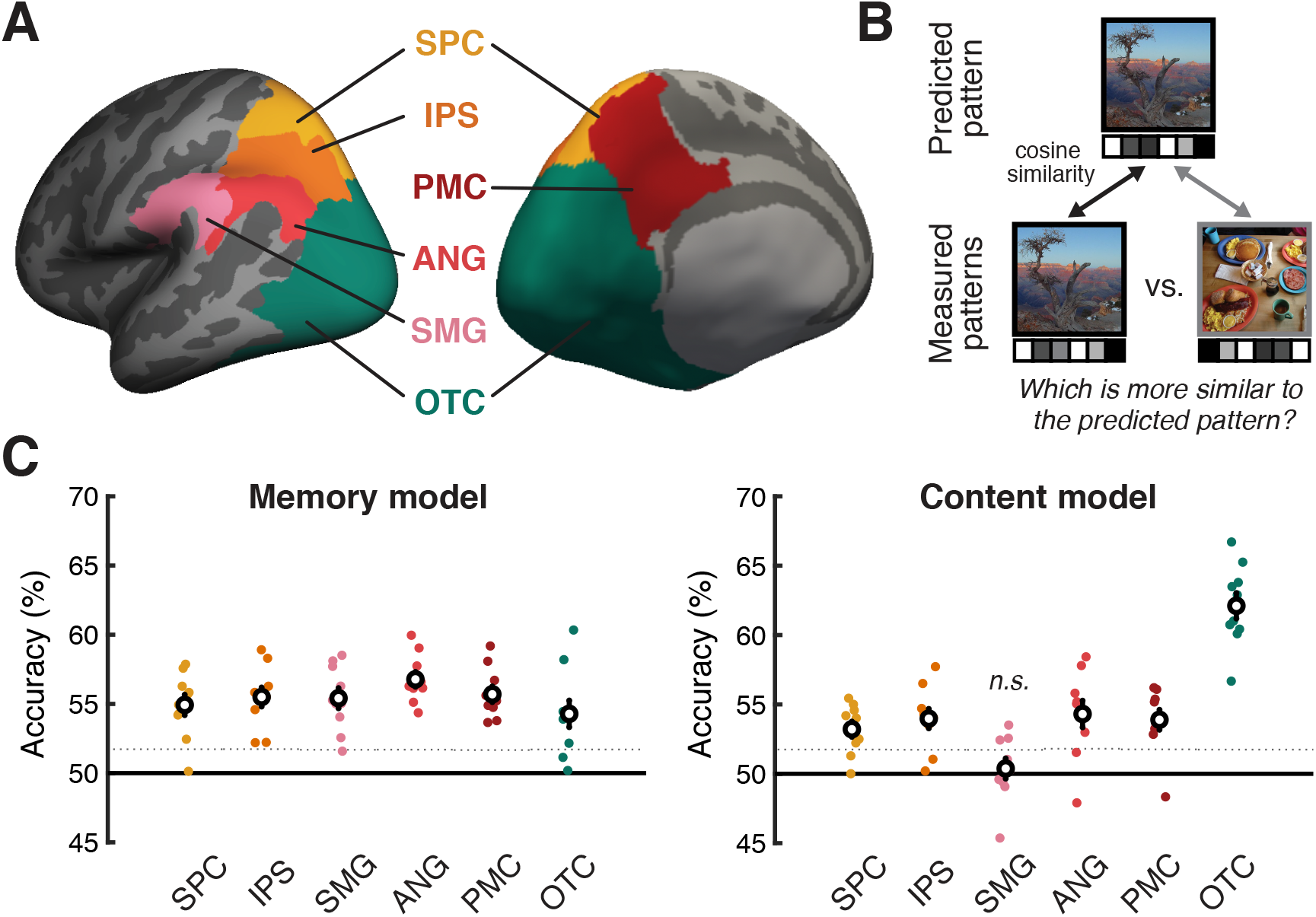
Prediction accuracy for the memory-based and content-based voxel-wise encoding models. (**A**) Regions of interest shown on the inflated cortical surface of the left hemisphere (left = lateral view; right = medial view). (**B**) Two-alternative-forced-choice (2AFC) test for assessing the trial-level accuracy with which the encoding models predicted the activation pattern within an entire region of interest. For each model (memory-based and content-based), we predicted the activation pattern within each region of interest for each image shown during the retrieval phase. The prediction was considered accurate when the predicted pattern was more similar to the actual (evoked) activation pattern than the pattern evoked by a different image. (**C**) 2AFC test accuracies of the memory model (left) and the content model (right). Colored dots indicate individual subjects. White circles indicate the mean across subjects. Error bars show SEM across subjects. Dotted horizontal lines show statistical significance thresholds (one-tailed, *p* < .05) defined from null distributions generated through permutation tests (averaged across all subjects). In **A** and **C**, SPC = superior parietal cortex, IPS = intraparietal sulcus, SMG = supramarginal gyrus, ANG = angular gyrus, PMC = posterior medial cortex, OTC = occipitotemporal cortex.

### General linear model analysis

To obtain trial-by-trial fMRI activation patterns for each subject and session, general linear model (GLM) analyses were performed using SPM12 (http://www.fil.ion.ucl.ac.uk/spm). The design matrix for each scanning run included 48 trial regressors convolved with the canonical hemodynamic response function. Six motion parameters and impulse responses representing volumes with unusually large motion (i.e., motion outliers detected using the function fsl_motion_outliers in FSL) were included as nuisance regressors. One-sample *t*-tests against a contrast value of 0 were performed to obtain trial-specific *t* statistic maps.

### Encoding model analysis

We created two separate voxel-wise encoding models: a memory-based encoding model and a content-based encoding model. The memory-based encoding model captured information about the episodic history of images and participants’ subjective memory judgments. The content-based encoding model captured information about the semantic content of images.

The memory-based encoding model included regressors representing the three levels of image repetition during the study phase (novel, repeated once, repeated three times) crossed by the four levels of recognition memory decisions (‘sure old,’ ‘likely old,’ ‘likely new,’ ‘sure new’). Separate regressors were generated for each trial type (e.g., novel images classified as ‘sure new’), with a value of one for the relevant trials and zero for the remaining trials. Additionally, while not a factor of interest—and not a factor that was controlled for when selecting stimuli— pre-experimental familiarity of images (or at least the potential for pre-experimental familiarity) was included as a regressor. Namely, trials in which the presented image contained a famous place or a famous person were assigned a value of one, and the remaining trials were assigned a value of zero. Thus, there were 13 regressors in total. We also built a separate memory-based encoding model excluding pre-experimental familiarity to assess the effect of the variable. All regressors were normalized (z-scored) across all trials included in the model.

The content-based encoding model was created using the human annotations collected from the online experiment (**Figure 1B-C**). Each online subject’s responses (words describing each image) were spell-checked and transformed into vectors of 300 numbers using Google’s pre-trained Word2Vec model. Words not included in the Word2Vec model were excluded from the analysis. For each image, the Word2Vec vectors were averaged across all words describing the image (responses were concatenated across online subjects) to generate a single vector per image. For dimensionality reduction (to reduce overfitting), a principal component analysis was performed on the Word2Vec vectors across all images used in the experiment. For each image, the first 30 principal component scores were selected to represent the image presented at each retrieval trial, and were used as the 30 regressors of the semantic encoding model. The first 30 principal components explained 70% of the variance among the vectorized descriptions.

For both the memory-based and content-based encoding models, we employed linear regression to predict the activation level of each voxel within an ROI for each retrieval trial from a single scanning session or both sessions. When a single scanning session was used, the trial-by-trial activation maps (*t* statistics) of an ROI were normalized (z-scored) across trials within the session. When both scanning sessions were used, the activation maps were first normalized across trials within each session, and then across all trials from both sessions. Trials in which subjects failed to make recognition memory responses were excluded from analyses. We used a leave-one-trial-out cross-validation method; we first generated the parameter estimates of the independent variables (regressors) using all but one trial included in the model, and used the parameter estimates to predict the activation of the left-out trial.

We tested the performance of the models using a two-alternative-forced-choice (2AFC) test method for which chance-level accuracy was 50% (**Figure 2B**) (Lee & Kuhl, 2016). Specifically, for each retrieval trial, we computed the cosine similarity between its predicted activation pattern and its measured (actual) activation pattern (same-image similarity). We also computed the cosine similarity between the predicted activation pattern of the trial and the measured activation pattern of every other trial (across-image similarity). We then separately compared the trial’s same-image similarity to each of the across-image similarity values. Thus, *N* – 1 2AFC tests were performed for each trial, where *N* is the total number of trials in the experiment. For each test, the prediction was considered accurate when same-image similarity was greater than across-image similarity. Thus, the accuracy for each trial was represented by the percentage of accurate predictions across all tests for that trial. These trial-level accuracy values were then averaged across trials (and sessions, where relevant) to generate a subject-specific prediction accuracy for each ROI. Within each ROI, and for each subject, we also identified individual voxels whose activation levels were significantly explained by the memory-based and/or content-based encoding models. To do this, we computed the Pearson correlation between the predicted time course of trial-by-trial activation levels and the measured activation time course for each voxel in an ROI (**Figure 3A**). We then generated a null distribution of correlations by randomly shuffling (1000 times) trial numbers and then re-computing the correlation between the predicted and measured activation time courses. The significance (one-tailed *p*-value) of the encoding model accuracy within the voxel was defined as the proportion of correlation values in the null distribution which were greater than or equal to the actual correlation between the predicted and measured time courses computed using the original (not shuffled) trial order.

**Figure 3.**
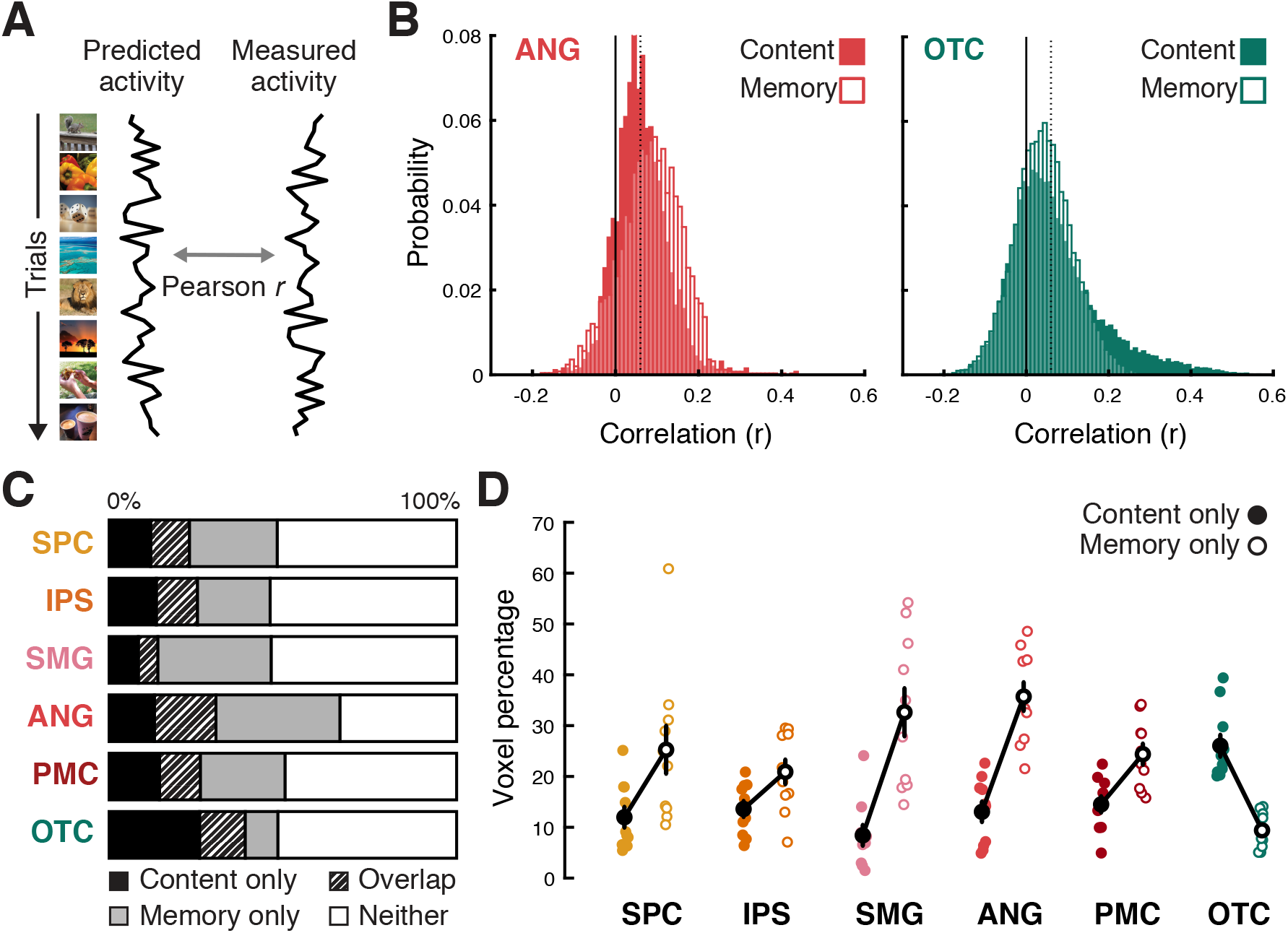
Voxel-specific encoding model accuracy and the distribution of different voxel types. (**A**) For each voxel, we measured the encoding model accuracy by computing the Pearson correlation between the predicted trial-by-trial activation time course (‘Predicted activity’; left) and the actual (evoked) time course (‘Measured activity’; right). (**B**) An example subject’s distribution of voxel-wise correlations between predicted and actual time courses in the angular gyrus (ANG; left) and the occipitotemporal cortex (OTC; right). Predictions were generated from either the content-based encoding model (filled bars) or the memory-based encoding model (unfilled bars). Voxels were considered to show significant effects for the content or memory models when their corresponding correlation values were greater than the statistical significance threshold (one-tailed, *p* < .05) defined from null distributions generated through permutation tests. Dotted vertical lines indicate the statistical threshold averaged across all voxels within each ROI (*r* = .06 for both models and regions). (**C**) Mean percentages of voxels within each region of interest that show significant effects of 1) the content model only (Content only), 2) the memory model only (Memory only), 3) both models (Overlap), and 4) neither model (Neither). (**D**) Percentages of voxels within each region of interest that were labeled as ‘content only’ or ‘memory only’. Colored filled/unfilled dots indicate individual subjects’ content/memory only voxel percentages, respectively. Black filled/unfilled dots indicate averages across subjects. In **C** and **D**, SPC = superior parietal cortex, IPS = intraparietal sulcus, SMG = supramarginal gyrus, ANG = angular gyrus, PMC = posterior medial cortex, OTC = occipitotemporal cortex.

For analyses related to encoding model accuracy and voxel distributions, we report the results obtained using data combined across both sessions. Results from single-session analyses were only used to independently select memory/content voxels that were then used for the content tuning analysis in a cross-validated manner (see below).

### Content tuning analysis

To characterize how individual brain regions combined episodic history and semantic content of images during retrieval, we generated content tuning functions for individual voxels within each ROI by measuring a given voxel’s response to different semantic categories of images. These tuning functions were separately generated for ‘hit’ and ‘correct rejection’ trials in order to assess whether the shape of the tuning function interacted with recognition memory.

We first categorized the image stimuli by applying K-medoids clustering (Lloyd, 1982) to the full Word2Vec vectors (not the dimensionally reduced data) for all images used in the experiment. Cosine distance was used as the distance metric to measure the similarity between each image vector and the cluster medoid. To select the number of clusters (k), we performed the clustering analysis using a range of k values (5 – 20), and selected k that maximized the number of clusters that had at least 80 images per cluster, after excluding images whose cosine distances to their corresponding cluster medoids were greater than .4. This resulted in 9 clusters (**Figure 4A**) roughly corresponding to the following semantic categories (as subjectively identified by the experimenters): humans (e.g., human faces, celebrities), animals, foods, architectural buildings, indoor scenes, urban outdoor scenes (e.g., streets, city landscapes), outdoor water scenes (e.g., beaches, lakes), mountains, events and activities (e.g., sports games). It is important to note that each image was assigned to only one of the 9 categories, despite the fact that images often combined elements from multiple categories (e.g., see images in **Figure 1**). Among the 1284 images used in the experiment, 146 images were excluded from the content tuning analysis as they were not strongly associated with any of the categories (i.e., cosine distance to the assigned cluster medoid > 0.4).

**Figure 4.**
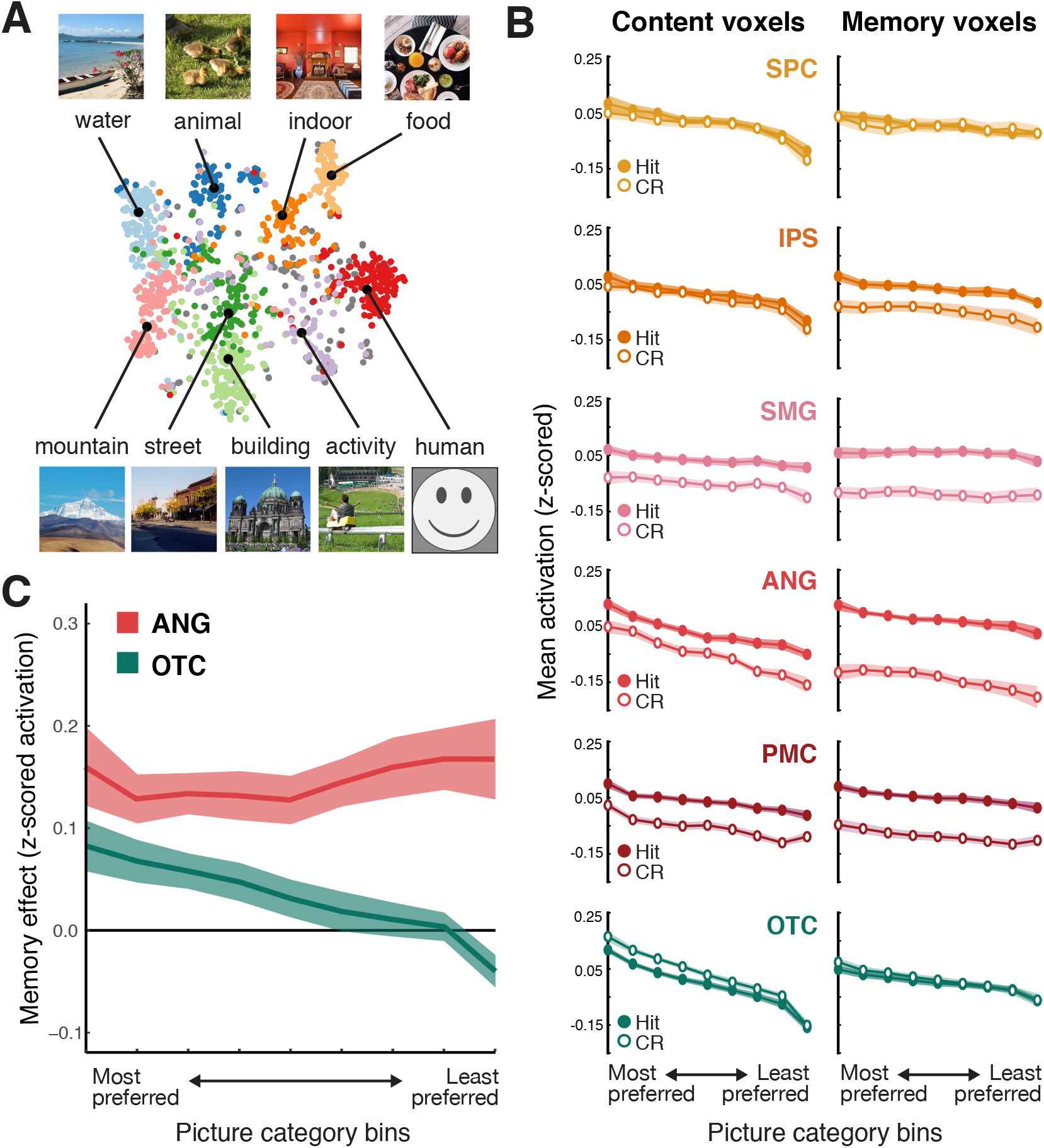
Content tuning analysis. (**A**) Nine semantic categories of image stimuli were identified by applying k-medoid clustering on the word embedding vectors describing the images. Each dot represents an image, located on a 2-dimensional space created by applying *t*-distributed stochastic neighbor embedding dimensionality reduction on the word embedding vectors of all images. Different colors represent different semantic categories. Each black dot and its associated picture show the medoid image of each category. The labels for each category (‘water’, ‘animal’, etc.) reflect subjective assessments (made by the experimenters) based on the clustering—these labels are purely descriptive. Note: the sample image for the human category is not an actual image from the experiment; actual images from the human category generally included human faces, which are withheld here due to publishing policies. (**B**) Content tuning functions for hit and correct rejection (CR) trials, separately for each ROI (row) and for content voxels (left) and memory voxels (right). Reliable content tuning is reflected by greater activation for ‘preferred’ semantic categories (left = most preferred category; right = least preferred category). (**C**) Memory effects (averaged across memory and content voxels) for angular gyrus (red) and occipitotemporal cortex (teal) as a function of voxels’ category preference. Note: for angular gyrus, memory effects were defined as z-scored activation for hit – correct rejections whereas for occipitotemporal cortex, memory effects were defined as z-scored activation for correct rejections – hits. Shaded areas indicate SEM across subjects.

We then generated tuning functions separately for each subject, voxel, and for the hit and correct rejection trials. Generating the tuning functions involved a two-step, cross-validated process where each session of the fMRI data was alternately used for each step (with results then averaged across the two cross-validation folds). The first step was to use data from one fMRI session to determine each voxel’s mean activation for each of the 9 semantic categories identified from the clustering analysis (see above). The mean activation per category was independently computed for the hit and correct rejection trials and data were *then* averaged across the hit and correct rejection trials (this ensured that hit and correct rejection trials had equal weight). For each voxel, the 9 semantic categories were then rank-ordered from the category that evoked the highest mean activation to the category that evoked the lowest mean activation. The second step was to test whether these category preferences, generated from half of the data, generalized to the held-out data (i.e., data from the other fMRI session). To do so, for each voxel we computed the mean activation for each rank-ordered semantic category using the held-out data. For a given voxel, its mean response for each of the rank-ordered semantic categories was defined as its content tuning function. To the extent that category preferences successfully generalized across fMRI sessions, this would be reflected by a negatively-sloped tuning function (i.e., decreasing activity for progressively ‘less preferred’ categories). Critically, content tuning functions were separately computed for hit trials and correct rejection trials by segregating trials from the held-out data according to memory status (hit, correct rejection) and then averaging across trials within each memory status (i.e., averaging the hit trials and averaging the correct rejection trials).

Finally, within each ROI we separately considered content tuning functions for (1) voxels that showed significant prediction effects for the memory-based encoding model (‘memory voxels’) and (2) voxels that showed significant prediction effects for the content-based encoding model (‘content voxels’). Significant prediction effects were defined as *p* < .05 (uncorrected) from the one-tailed permutation tests described in *Encoding Model Analyses*. The rationale for comparing tuning functions for ‘memory voxels’ vs. ‘content voxels’ was to test whether some voxels preferentially—or even selectively—carried content information or memory information. Critically, however, the encoding models used for voxel selection were based on only half of the data (the same half of the data used to identify the content tuning preferences). Thus, the voxel selection procedures (i.e., how the voxel groups were defined) and the category preference procedures (i.e., how tuning functions were defined) were based on fMRI data that was entirely independent from the critical test data. Again, this ensured that there was no circularity to these analyses (Kriegeskorte et al., 2009).

In summary, our procedure for generating content tuning functions allowed us to test whether the relative content preferences of a voxel—which were defined using half of the data— generalized to independent, held-out data. This cross-validated procedure critically ensured that there was no circularity in how the content tuning functions were generated. Importantly, while tuning functions were always generated in a voxel-specific manner and individual voxels were likely to have different category preferences, our approach readily allowed for tuning functions to be averaged across voxels within an ROI. That is, our approach tested the degree to which each voxel’s category preferences were preserved (across sessions), regardless of whether individual voxels had similar category preferences.

### Statistical tests

To test the accuracy of our 2AFC test, where chance performance would correspond to 50% accuracy, we used two-tailed, one-sample *t*-tests to compared observed accuracy to 50%. For direct comparison between pairs of ROIs or conditions, we performed two-tailed, paired-samples *t*-tests. For comparisons involving more than two ROIs/conditions or for testing interactions between different factors, we used repeated-measures ANOVAs, with subject number as the error factor. Within the ANVOAs, linear contrast analyses were used to test for linear trends along the rank ordered category preference bins in the content tuning analysis.

To test the significance of encoding model accuracy within each individual subject, we performed nonparametric permutation tests. For each subject, we generated the null distribution of accuracy by randomly shuffling the trial numbers and then computing the subject-specific accuracy using the same 2AFC test method as described above in *Encoding model analysis* (number of iterations = 1000). The significance (one-tailed *p*-value) of subject-specific model accuracy was defined as the proportion of accuracy values in the null distribution which were greater than or equal to the actual accuracy computed using the original trial order.

## RESULTS

### Behavioral results

All behavioral data were first averaged across sessions and then across subjects. Data are reported as mean (M) ± standard deviation (SD). During the study phase (which occurred outside the fMRI scanner), the mean percentages of trials receiving each pleasantness rating were as follows: ‘like’ = 55.28% ± 19.35%, ‘dislike’ = 11.89% ± 11.13%, ‘neutral’ = 30.70% ± 18.67%, no response = 2.12% ± 5.89%. The mean percentage of images with a consistent response across all three presentations was 80.51% ± 5.94%. Because pleasantness ratings were only included as an incidental encoding task, these data are not considered further. During the scanned recognition memory task, the mean hit rate was 88.51% ± 8.93% and the mean false alarm rate was 9.82% ± 7.77%. Mean sensitivity (measured by d’) was 2.77 ± .80 (range: 1.37 – 4.02), which was significantly above chance (*t*_9_ = 10.97, *p* < 0.0001). Sensitivity did not differ between Sessions 1 and 2 (*t*_9_ = 0.29 *p* = .779). Sensitivity was significantly higher for images that were presented three times in the study phase compared to those presented once (d’ = 3.37 ± .38 vs. 2.47 ± .47, respectively; *t*_9_ = 8.41, *p* < 0.00007).

### Memory-based encoding model

The memory-based encoding model attempted to predict the activity patterns evoked by each scene image based on three variables: the number of repetitions in the study phase, response during the recognition memory task, and pre-experimental familiarity (see Methods for details). Prediction accuracy was assessed for each image by using cosine similarity to compare the predicted activity pattern to (a) the activity pattern evoked by that image (same-image similarity), and (b) the activity pattern evoked by other images (across-image similarity; see Methods). If same-image similarity exceeded across-image similarity for a given comparison, this was considered an accurate prediction. Using this method, chance accuracy corresponds to 50%.

Accuracy was above chance for each of the ROIs, as assessed by one-sample *t*-tests (SPC: 54.93 ± 2.21%, *t*_9_ = 6.68, *p* = 0.0001; IPS: 55.50 ± 2.07%, *t*_9_ = 7.99, *p* < 0.0001; SMG: 55.41 ± 2.19%, *t*_9_ = 7.42, *p* < 0.0001; ANG: 56.78 ± 1.61%, *t*_9_ = 12.60, *p* < 0.0001; PMC: 55.70 ± 1.70% *t*_9_ = 10.06, *p* < 0.0001; OTC: 54.28 ± 2.89%, *t*_9_ = 4.45, *p* = 0.002; **Figure 2C**). However, accuracy also significantly varied across the ROIs (repeated-measures ANOVA: *F*_5,45_ = 5.04, p = 0.001), and was numerically highest in ANG and lowest in OTC. A direct contrast between ANG and OTC revealed significantly greater accuracy in ANG (paired-samples *t*-test: *t*_9_ = 4.67, *p* = 0.001). Considering individual subjects, accuracy was above chance, as determined by subject-specific permutation tests (one-tailed, *p* < .05; see Methods), for every subject (10/10) for ANG, IPS, and PMC; for 9/10 subjects for SPC and SMG; and for 8/10 subjects for OTC.

We also tested whether excluding the pre-experimental familiarity variable (whether or not scenes had ‘famous’ content) had any impact on the memory-based encoding model accuracy. Removing this variable from the model resulted in slightly but significantly lower accuracies in ANG (−0.28 ± 0.29%, *t*_9_ = 3.04, *p* = 0.014) and PMC (–0.28 ± 0.35%, *t*_9_ = 2.51, *p* = 0.033), but no significant difference for other parietal ROIs (SPC: –0.05 ± 1.0%, *t*_9_ = 0.14, *p* = 0.89; IPS: –0.48 ± 0.80%, *t*_9_ = 1.88, *p* = 0.09; SMG: –0.13 ± 0.52%, *t*_9_ = 0.79, *p* = 0.45). For OTC, however, accuracy significantly increased when pre-experimental familiarity was excluded (2.81 ± 1.68%, *t*_9_ = −5.27, *p* = 0.0005). These data indicate that pre-experimental familiarity positively contributed to prediction accuracy only for ANG and PMC, and very modestly for these regions. Ultimately, we do not consider this variable in more detail given the fact that the number of ‘famous’ vs. ‘non-famous’ scenes was not balanced across subjects, runs, or repetition conditions. That said, our rationale for including the variable in the memory-based encoding model was that it does reflect a form of memory for an image and might therefore explain meaningful variance.

### Content-based encoding model

The content-based encoding model attempted to predict the activity pattern evoked by a scene image based on the 30 principal components that represented the content of the image (see Methods). Prediction accuracy was assessed using the same procedures as for the memory-based encoding model (i.e., by comparing same-image similarity to across-image similarity). Accuracy was above chance for each of the ROIs except SMG, as assessed by one-sample *t*-tests (SPC: 53.2 ± 1.65%, *t*_9_ = 5.84, *p* = 0.0002; IPS: 54.0 ± 2.10%, *t*_9_ = 5.68, *p* = 0.0003; SMG: 50.4 ± 2.18%, *t*_9_ = 0.52, *p* = 0.615; ANG: 54.3 ± 2.88%, *t*_9_ = 4.48, *p* = 0.0015; PMC: 53.89 ± 2.16%, *t*_9_ = 5.42, *p* = 0.0004; OTC: 62.1 ± 2.75%, *t*_9_ = 13.19, *p* < 0.0001). Accuracy markedly varied across the ROIs (repeated-measures ANOVA: *F*_5,45_ = 58.08, *p* < 0.0001; **Figure 2C**), with OTC exhibiting the highest accuracy. While ANG exhibited the highest accuracy (numerically) among parietal ROIs, ANG accuracy was significantly lower than accuracy in OTC (paired-samples *t*-test: *t*_9_ = 16.34, *p* < 0.0001). Considering individual subjects, accuracy was above chance, as determined by subject-specific permutation tests (one-tailed *p* < .05), for every subject (10/10) for OTC; for 9/10 subjects for PMC; for 8/10 subjects for SPC, IPS, and ANG, and for 3/10 subjects for SMG.

To test whether the content model and the memory model were differentially predictive of activity patterns across regions of interest, we performed a repeated measures ANOVA with factors of ROI (all ROIs) and encoding model type (memory, content). The interaction between ROI and model type was highly significant (*F*_6,54_ = 49.364, *p* < 0.0001). This interaction was largely driven by the fact that OTC was associated with higher accuracy than the parietal ROIs for the content model, but lower accuracy than the parietal ROIs for the memory model. There was also a significant interaction when directly comparing ANG vs. OTC (*F*_1,9_ = 191.34, *p* < 0.0001).

### Distribution of memory and content voxels

We next assessed the percentage of voxels within each ROI that exhibited significant effects for each encoding model (the memory-based and content-based models; see Methods, **Figure 3B-D**). For this analysis, we used a liberal threshold (*p* < 0.05) as the goal was only to assess the relative distribution of these voxels across ROIs. Within each ROI, each voxel was labeled according to one of four categories: content only, memory only, overlap (i.e., both), or neither (**Figure 3C**). As a first step, we compared the mean percentage of content versus memory voxels (excluding overlap voxels) across ROIs (**Figure 3D**). An ANOVA with factors of voxel type (content, memory) and ROI revealed a significant main effect of ROI (*F*_5,45_ = 4.86, *p* = 0.001), a significant main effect of voxel type (*F*_1,9_ = 12.57, *p* = 0.006), and a significant interaction between ROI and voxel type (*F*_5,45_ = 14.216, *p* < 0.0001). The main effect of ROI was primarily driven by the relatively high percentage of significant voxels (content *and* memory) in angular gyrus. The main effect of voxel type reflected an overall higher percentage of memory voxels (*M* = 24.71 ± 8.50%) compared to content voxels (*M* = 14.60 ± 5.47%). However, the significant interaction reflected the fact that whereas parietal ROIs exhibited relatively more memory voxels than content voxels, OTC exhibited the opposite pattern. Paired-samples *t*-tests applied to the individual ROIs revealed an effect/trend toward a higher percentage of memory voxels compared to content voxels for each of the parietal ROIs (SPC: 25.25 ± 4.77% vs. 11.98 ± 2.09%, *t_9_* = 2.24, *p* = 0.052; IPS: 20.90 ± 2.47% vs. 13.60 ± 1.63%, *t_9_* = 2.01, *p* = 0.076; SMG: 32.63 ± 4.79% vs. 8.41 ± 2.10%,, *t_9_* = 3.98, *p* = 0.0032; ANG: 35.71 ± 2.88% vs. 13.04 ± 2.07%, *t_9_* = 5.08, *p* = 0.0007; PMC: 24.38 ± 2.14% vs. 14.50 ± 1.66%, *t_9_* = 2.99, *p* = 0.015); for OTC, however, there was a significantly lower percentage of memory voxels than content voxels (9.40 ± 1.11% vs. 26.04 ± 2.21%, *t_9_* = −5.48, *p* = 0.0004). A direct comparison between ANG and OTC revealed a significant interaction between ROI and voxel type (*F*_1,9_ = 40.18, *p* = 0.0001) again reflecting the relative bias toward memory voxels in ANG and content voxels in OTC.

We also compared the total percentage of significant voxels (content + memory + overlap) across the ROIs. A repeated-measures ANOVA revealed a significant main effect of ROI (*F*_5,45_ = 8.93, *p* < 0.0001) with ANG again containing the highest percentage of total significant voxels (**Figure 3C**). Similarly, considering the percentage of overlap voxels alone, there was a significant main effect across ROIs (*F*_5,45_ = 5.71, *p* = 0.0003), with ANG containing the highest percentage of overlap voxels and SMG containing the lowest. Direct contrasts (paired-samples *t*-tests) between ANG and OTC revealed a higher total percentage of significant voxels (content + memory + overlap) in ANG compared to OTC (ANG: 66.46 ± 14.85%; OTC: 48.59 ± 9.16%; *t*_9_ = 5.05, *p* = 0.0007) but no significant difference in the percentage of overlap voxels (ANG: 17.70 ± 3.45%; OTC: 13.14 ± 2.37%; *t*_9_ = 1.09, *p* = 0.29).

### Content tuning as a function of recognition memory decisions

The preceding analyses indicate that content information and memory information were broadly distributed across parietal cortex and OTC, while also highlighting differences in how information was distributed across regions. To better characterize how individual brain regions *combined* memory and content information, we conducted a complementary series of analyses in which we generated ‘content tuning functions’ for each ROI. Critically, separate tuning functions were generated for *hits* (old scenes endorsed as ‘old’) and *correct rejections* (new scenes endorsed as ‘new’) in order to test whether content tuning differed as a function of recognition memory status (for details, see Methods). To generate these tuning functions, we first used k-medoids clustering applied to scene image annotations to group the scene images into 9 semantic categories (see Methods; **Figure 4A**). For every voxel within each ROI, half of the fMRI data (i.e., data from one fMRI session) were used in a cross-validated manner to rank-order the 9 categories according to the voxel’s ‘preference’ (i.e., its relative activation to images from each category). These voxel-specific preferences were then tested for generalization in the held-out data (i.e., data from the other session). Successful generalization would be evidenced by voxels displaying the same relative profile of activation across categories (i.e., the same ‘tuning’) in the held-out data. Specifically, we tested for a linear trend in activation as a function of category preference (i.e., that activation decreased from the ‘most’ to ‘least’ preferred categories). Finally, within each ROI, we compared tuning functions for: (1) voxels that exhibited significant content effects (content voxels) and (2) voxels that exhibited significant memory effects (memory voxels), as defined based on results from independent encoding model analyses (see Methods). The rationale for separately considering content voxels and memory voxels (and for excluding the overlap voxels) is that it provides another way for assessing the separability (or inseparability) of content and memory information. Namely, if voxels specifically selected for exhibiting *either* content or memory effects (in one half of the data) nonetheless express the other form of information (in the held-out half of data), this would provide evidence that these two forms of information are highly overlapping (or, put another way, difficult to separate).

For each ROI, we generated a separate repeated-measures ANOVA with factors of semantic category preference (the 9 rank-ordered category preference bins), recognition memory status (hit, correct rejection), and voxel type (content, memory). We first tested for main effects of semantic category preference (combining across the content and memory voxels): that is, whether the semantic tuning preferences identified from half of the data (one session) generalized to the held-out data (the other session). Indeed, for each ROI, there was a significant linear trend as a function of category preference (see *Category Effects* in **Table 1**). These linear trends indicate that category preferences were preserved across independent sessions and validate our approach for generating semantic tuning functions. Notably, the linear trends of semantic category preference interacted with voxel type (content vs. memory voxels) for all ROIs (**Table 1**). In each case, this interaction reflected a stronger linear trend (‘sharper’ tuning) for content voxels than memory voxels (see **Figure 4B**).

**Table 1.** Analysis of content tuning functions. For each region of interest (ROI) separate ANOVAs were applied to the content tuning functions to test for (1) linear effects of semantic category preference (across the 9 semantic categories; Category Effects); (2) interactions between Category Effects and voxel type (memory voxels vs. content voxels; Category * Voxel Type); (3) main effects of memory (hits vs. correct rejections; Memory Effects); and (4) interactions between Memory Effects and voxel type (Memory * Voxel Type). Notes: Category Effects and Memory Effects included voxel type (memory voxels vs. content voxels) as a factor, but overlap voxels were excluded; for all ANOVAs, degrees of freedom = 1,9.

| ROI | Category Effects<br>(content tuning) |  | Category *<br>Voxel Type |  | Memory Effects<br>(hit vs. CR) |  | Memory *<br>Voxel Type |  |
| --- | --- | --- | --- | --- | --- | --- | --- | --- |
|  | *F* | *p* | *F* | *p* | *F* | *p* | *F* | *p* |
| ANG | 319.38 | < 0.0001 | 37.24 | < 0.0001 | 90.78 | < 0.0001 | 67.89 | < 0.0001 |
| IPS | 248.96 | < 0.0001 | 19.93 | 0.002 | 4.65 | 0.06 | 11.83 | 0.007 |
| SMG | 38.16 | < 0.0001 | 10.72 | 0.02 | 32.68 | 0.0003 | 7.25 | 0.03 |
| SPC | 164.31 | < 0.0001 | 32.72 | < 0.0001 | 0.16 | 0.70 | 0.32 | 0.59 |
| PMC | 228.75 | < 0.0001 | 11.72 | 0.0007 | 112.56 | < 0.0001 | 8.07 | 0.02 |
| OTC | 1762.1 | < 0.0001 | 300.43 | < 0.0001 | 1.77 | 0.22 | 7.98 | 0.02 |

We next tested for main effects of memory (hit vs. correct rejection). Significant main effects of memory were observed in SMG, ANG, and PMC, but not in IPS, SPC or OTC (see *Memory Effects* in **Table 1**). Interactions between memory and voxel type were present in all of the ROIs except SPC (SPC: *F*_1,9_ = 0.32, *p* = 0.59, IPS: *F*_1,9_ = 11.84, *p* = 0.007 SMG: *F*_1,9_ = 7.25, *p* = 0.03, ANG: *F*_1,9_ = 67.89, *p* < 0.0001, PMC: *F*_1,9_ = 8.07, *p* = 0.02; OTC: *F*_1,9_ = 7.98, *p* = 0.02; **Table 1**). Interestingly, however, while this interaction reflected a stronger effect of memory (hit > correct rejection) for memory voxels compared to content voxels in the parietal ROIs, OTC exhibited precisely the opposite effect, with a stronger effect of memory (in the direction of hits < correct rejections) for content voxels compared to memory voxels (**Figure 4B**). The fact that memory effects in OTC were stronger for content voxels than memory voxels suggests that memory effects in OTC were fundamentally related to a voxel’s content sensitivity. This finding is consistent with prior evidence that repetition suppression signals in OTC are content dependent (Grill-Spector et al., 2006).

Finally, and critically, we tested whether the shape of the content tuning functions differed as a function of recognition memory status (hit, correct rejection). An ANOVA that again included factors of category preference (the 9 semantic categories), recognition memory status (hit, correct rejection), and voxel type (content voxels, memory voxels) revealed a significant interaction between category preference (the linear trend across category preference bins) and memory (hit, correct rejection) for OTC (repeated measures ANOVA: *F*_8,72_ = 13.37, *p* = 0.0003), but not for any of the parietal ROIs (SPC: *F*_8,72_ = 1.87, *p* = 0.17, IPS: *F*_8,72_ = 0.001, *p* = 0.97 SMG: *F*_8,72_ = 0.02, *p* = 0.90, ANG: *F*_8,72_ = 3.47, *p* = 0.06, PMC: *F_8_*_,72_ = 0.22, *p* = 0.64). (Note: none of the ROIs exhibited a three-way interaction between semantic category, memory status and voxel type: all *p*’s > 0.32). For OTC, the interaction between semantic category and memory status reflected a relatively stronger effect of memory (correct rejection > hit) for ‘preferred’ semantic categories.

To formally contrast the relationships between memory effects and content effects in ANG vs. OTC, we computed, for each ROI and averaging across content and memory voxels, the size of the memory effect (the difference between hits vs. correct rejections) for each category preference bin. Importantly, however, we defined memory effects in ANG as *hit – correct rejection*, whereas for OTC memory effects were defined as *correct rejection – hit*. The rationale for this different definition across ROIs is that here, and in numerous prior studies, memory effects in ANG are reflected by increased activation for ‘old’ items (repetition enhancement) (Cabeza et al., 2008; Rugg & King, 2018; Sestieri et al., 2017; Wagner et al., 2005) whereas memory effects in OTC are reflected by decreased activation for ‘old’ items (repetition suppression) (Grill-Spector et al., 2006; Miller et al., 1991). Thus, this allowed us to compare the absolute magnitude of memory effects across ANG and OTC. Indeed, there was a significant interaction between ROI (ANG, OTC) and category preference (the linear trend) (*F*_1,9_ = 6.605, *p* = 0.01; **Figure 4C**), confirming that memory effects in ANG and OTC were differentially sensitive to category preference. As described above, memory effects in OTC were relatively stronger for more ‘preferred’ semantic categories. In contrast, memory effects in ANG were robust and generally consistent across category preference bins. Thus, despite the fact that ANG and OTC each contained information about memory and content, these regions combined these forms of information in distinct ways.

## DISCUSSION

Here, we used voxel-wise encoding models and content tuning functions to characterize content representations of natural scene images in parietal and occipitotemporal cortices during a recognition memory task. We show that memory- and content-related signals are robustly distributed and highly overlapping within parietal cortex—particularly within the angular gyrus. While these two forms of information were expressed within common voxels in angular gyrus, they were statistically independent: content tuning did not interact with memory effects. In contrast, memory effects in occipitotemporal cortex (OTC) were preferentially carried by content-sensitive voxels and the magnitude of these effects was dependent on the degree to which a given OTC voxel ‘preferred’ the content of a remembered stimulus. These findings provide new insight into how the brain combines content and memory information.

Our findings are consistent with numerous fMRI studies of human memory showing that content information can be decoded from parietal cortex (Bird et al., 2015; Bonnici et al., 2016; Favila et al., 2018; Kuhl & Chun, 2014; Lee et al., 2016, 2018). However, our combination of semantic (content-based) encoding models (Huth et al., 2016; Pereira et al., 2018) and content tuning functions provides a richer and more rigorous characterization of content representations than typical decoding measures. In particular, the content representations measured here cannot be explained as category or stimulus labels that are adaptively generated to satisfy task demands (Toth & Assad, 2002). Rather, the encoding model represents content as continuous weights across a diverse set of features (Bone et al., 2020; Lee & Kuhl, 2016). Moreover, the models were trained on images that were distinct from test images, avoiding the possibility that the model learned stimulus-specific labels. For the tuning functions, although we grouped images into 9 semantic categories, this was done for dimensionality reduction and these categories were not behaviorally-relevant to subjects. Thus, our findings provide some of the most compelling evidence to date that parietal regions involved in episodic memory also encode rich and multidimensional content information (Bird et al., 2015; Bone et al., 2020; Huth et al., 2016).

By generating separate encoding models for memory- and content-related information, we were able to compare the relative sensitivity of parietal and occipitotemporal regions to each type of information. Not surprisingly, content effects were stronger in OTC than in parietal cortex (**Figure 2C**, **3B-D**). Within parietal cortex, however, there were qualitative differences across sub-regions. For example, while supramarginal gyrus and angular gyrus were both characterized by relatively strong memory effects (**Figure 2C, 3C-D**), content effects were more apparent in angular gyrus than in supramarginal gyrus (**Figure 2C**). Thus, our findings provide a unique characterization of functional heterogeneity across parietal regions (Hutchinson et al., 2014; Sestieri et al., 2017). In particular, our findings support the idea that, among parietal regions, angular gyrus was uniquely sensitive to the combination of memory and content information (**Figure 3C**) (Bonnici et al., 2016; Humphreys et al., 2021; Kuhl & Chun, 2014).

For our tuning function analyses, we sought to more precisely determine whether content representations changed as a function of recognition memory (Woolnough et al., 2020). Critically, we first *independently identified* subject-specific voxels from the encoding models based on whether they exhibited memory or content effects. Notably, we excluded ‘overlap voxels’ in order to test whether specific populations of voxels *selectively* expressed either memory or content information. Within angular gyrus, memory effects (hit > correct rejection) were, not surprisingly, stronger in memory voxels than content voxels and, conversely, content tuning was stronger (sharper tuning) in content voxels than memory voxels. In other words, angular gyrus contained voxels that preferentially expressed either memory or content effects. Critically, however, content voxels still expressed memory effects and memory voxels still expressed content effects. Thus, even when we deliberately attempted to select voxels in angular gyrus that *only* expressed one type of information based on half of the data, these voxels nonetheless strongly expressed *both* forms of information in the held-out data. Thus, memory and content effects were not merely overlapping in angular gyrus (Bonnici et al., 2016; Kuhl & Chun, 2014; Ramanan et al., 2018; Rugg & King, 2018; Shimamura, 2011)—they were difficult to segregate.

Our second main finding from the tuning function analyses was that the *shape* of content tuning in angular gyrus was unaffected by recognition memory success. Specifically, when contrasting tuning functions for hits versus correct rejections, there was a shift in the tuning functions (hit > correct rejection), but the tuning functions were parallel. Put another way, recognition-related increases were unrelated to a voxel’s preference for the content of the recognized image. Thus, although individual voxels in angular gyrus were reliably tuned to different types of content and these same voxels also strongly reflected recognition memory success, content representations were invariant to recognition memory success.

Importantly, the tuning function results in angular gyrus statistically contrasted with OTC. First, memory effects in OTC tuning functions (hit < correct rejection) were stronger for *content voxels* than for memory voxels. While counterintuitive, this suggests that memory effects in OTC were secondary to, or derived from, content representations. Indeed, the number of content voxels in OTC was also much higher than the number of memory voxels (**Figure 3C**)—thus, selecting voxels on the basis of content sensitivity was a more effective form of feature selection. Second, there was a statistical interaction between memory effects and content tuning in OTC. Namely, OTC memory effects were stronger for voxels that ‘preferred’ the content of the remembered image. Importantly, this interaction in OTC statistically differed from the relative independence of content and memory effects in angular gyrus (and other parietal regions) (**Figure 4B-C**). Thus, in contrast to angular gyrus, memory effects in OTC scaled with the degree to which voxels preferred the content of a recognized image (Grill-Spector et al., 2006).

Collectively, our findings are consistent with theoretical accounts which argue that angular gyrus functions as a convergence zone for multiple sources of information during memory retrieval (Ramanan et al., 2018; Seghier, 2013; Tibon et al., 2019) and that angular gyrus jointly contributes to both semantic and episodic memory (Humphreys et al., 2021). While our findings cannot adjudicate between all of the competing theories of how the angular gyrus contributes to memory (Cabeza et al., 2008; Vilberg & Rugg, 2008; Wagner et al., 2005), a unique conclusion we can draw is that univariate increases in angular gyrus during successful recognition were not *driven by* the representation of recognized content. Instead, recognition effects may be better characterized as a broadband signal that rides on top of feature-specific channels that are tuned to different types of content. One interesting possibility is that recognition memory signals in angular gyrus could reflect a global bias toward internal processing that is induced by successful recognition (Honey et al., 2017). However, it is important to emphasize that our findings are based on a recognition memory paradigm and, therefore, may not generalize to other forms of memory (cued or free recall) where to-be-remembered content must be internally-generated as opposed to being perceptually available. Interestingly, there is evidence that content representations in angular gyrus are actually stronger during cued recall than during perception (Favila et al., 2018; Long & Kuhl, 2021; Xiao et al., 2017). It would therefore be informative to apply the analyses used here to more thoroughly compare content representations during recall versus perception. Another issue that is beyond the scope of the current study is the degree to which the effects reported here depend on the subjective experience of remembering versus the objective experience of stimulus repetition. A compelling body of evidence indicates that the angular gyrus is involved in (Hutchinson et al., 2014; Kuhl & Chun, 2014; Ramanan et al., 2018) and even necessary for (Simons et al., 2010; Tibon et al., 2019; Yazar et al., 2014; Zou & Kwok, 2022) the subjective experience of remembering. In contrast, memory effects in OTC may be more closely related to objective effects of stimulus repetition (Sayres & Grill-Spector, 2006; Ward et al., 2013). Here, we were not able to tease these apart because the proportion of ‘miss’ trials (objectively ‘old’ but subjectively ‘new’ trials) was very low. Finally, it would also be informative to consider the relative timing of content and memory representations across angular gyrus and OTC (Staresina & Wimber, 2019). While difficult to address with fMRI, intracranial electrophysiological measures have the potential to provide unique insight into these dynamics (Gonzalez et al., 2015).

In summary, our findings provide new insight into how the brain combines information about ‘what’ is being remembered with information about ‘whether’ something is being remembered. By directly contrasting memory and content effects across different brain regions, we show that there are multiple ways in which the brain combines these two forms of information. Our findings will hopefully inform and constrain theoretical accounts of parietal contributions to memory and inspire new, targeted research studies that further characterize how content and memory signals are combined in the brain.

## Author contributions

H.L., S.C.S., and B.A.K designed the experiment; H.L., S.C.S., and P.A.K. collected and analyzed data; H.L., P.A.K., J.B.H., and B.A.K. wrote and edited the manuscript.

## Acknowledgments

The research reported here was supported by NSF CAREER Award BCS-1752921 and NIH-NINDS 1R01 NS107727 to B.A.K. The authors have no competing financial interests to declare.

